# Uncertainty in the mating strategy causes bias and inaccuracy in estimates of genetic parameters in honeybees

**DOI:** 10.1101/2023.05.22.541688

**Authors:** Tristan Kistler, Evert W. Brascamp, Benjamin Basso, Piter Bijma, Florence Phocas

## Abstract

**Background:** With the increased number of honeybee breeding plans worldwide, records from queens with diversified mating strategies need to be considered. Breeding queens might be inseminated with drones produced by a single drone-producing queen (DPQ), or by a group of sister-DPQs. Often, only the dam of DPQ(s) is reported in the pedigree. Furthermore, datasets might include colony phenotypes from DPQs that were open mated in different locations. Using simulation, we investigated the impact of the mating strategy on estimates of genetic parameters and breeding values, when the DPQs were treated in different ways in the statistical evaluation model. We quantify the bias and standard error of estimates when breeding queens are mated to a single or a group of DPQs, assuming that this information is either known or not. We also investigated two alternative strategies to accommodate phenotypes of open-mated DPQs in the genetic evaluation, adding either a dummy pseudo sire in the pedigree, or a non-genetic effect to the statistical evaluation model to account for the origin of the DPQs’ mates.

**Results:** When breeding queens were inseminated with semen from drones of a single DPQ and this was known, estimates of genetic parameters and genetic trends were more precise. If they were inseminated using drones from a single or a group of DPQs, and this information was not known, erroneous assumptions led to considerable bias in the estimates. For colony phenotypes of open-mated DPQs, adding a dummy pseudo sire in the pedigree for each mating location led to considerable overestimation of genetic variances, while correcting for the mating area by adding a non-genetic effect in the evaluation model gave unbiased estimates.

**Conclusions:** Knowing only the dam of the DPQ(s) in the mating may lead to erroneous assumptions on how DPQs were used and cause severe biases in estimates of genetic parameters and genetic trends. Therefore, keeping track in the pedigree of which DPQ(s), and not only which dam of DPQ(s) are used, is recommended. Records from DPQ colonies with queens open mated to a heterogeneous drone population can be integrated by adding non-genetic effects to the statistical evaluation model.

## Background

Although mating control is essential to genetic improvement in a honeybee breeding program [1], its practical application is not straightforward, due to queens’ behavioral and anatomical peculiarities. Indeed, natural mating occurs during flight, typically at a few tens of meters high, where drones and young queens from several kilometers around gather together [2]. In order to control the genetic origin of mates in a breeding program, virgin queens and mature drones can be geographically isolated on mating stations, or alternatively, artificial insemination can be used.

At mating stations used in selective breeding, usually a group of sister drone-producing queens (DPQs) descending from a single dam is used to produce all the drones of a mating station. This group is referred to as a pseudo sire (PS) and is registered in the pedigree. To ensure high mating success, the group is usually composed of around four to a dozen of DPQs (personal communication from the French royal jelly producers: GPGR, and island mating in the Beebreed dataset [3]). Virgin queens brought to the mating station are then mated by the drones on the mating station. We call this PS mating.

The other alternative, artificial insemination, allows for greater mating control. In particular, drones used to mate a virgin queen can be taken from a PS composed of very few sister-DPQs or even from a unique DPQ (for example in [4]). In the latter case, we call it single sire (SS) mating. Compared to PS mating, SS mating generates more related female offspring in the colony. When properly accounted for in the pedigree, SS mating should enable estimations of genetic parameters and breeding values with lower standard errors than PS mating for which the precise origin of drones cannot be distinguished among the sister-DPQs and has to be probabilistically derived [5,6]. However, when artificial insemination is used, honeybee breeders often only record the dam of the DPQ(s) and provide no information about the number of sister-DPQs, even when only one DPQ is involved.

In addition, selective breeding programs also commonly include open mating, in which virgin queens are allowed to mate in unrestricted geographic areas (for example in Italy [7]). In particular, DPQs are often open mated (for example in France [8]), because this reduces managing costs. If the contribution of DPQs to the breeding population is limited to producing drones, then their mates do not affect the genetic evaluation because drones are haploid individuals born from unfertilized eggs, and thus do not carry genes from their dam’s mate. With artificial insemination, however, these DPQs are often phenotyped and are used for drone production after phenotypic selection. In that case, the mates of DPQs affect the genetic evaluation as they are the sires of the workers of the DPQ colonies, and the record of the colonies are used in the genetic evaluation.

For most traits, the phenotypes of a colony are supposed to be affected by both the queen and its worker group. The colony performance should therefore be partitioned into two genetic effects, a worker (or direct) genetic effect expressed by the worker group, and a queen (or maternal) genetic effect expressed by the workers’ dam, in addition to environmental effects [9,10]. Hence, instead of a single genetic variance, three genetic parameters need to be estimated: the variances of queen and worker effects, and their covariance. A reliable estimation of genetic parameters requires data and pedigree records from a large population of genetically well-connected apiaries. Unfortunately, most honeybee breeding programs use small nucleus populations of ten to a few tens of breeding queens [11–13,8]. In addition, queens only mate before their first egg lay, and will normally never mate afterwards. This adds to the difficulty of disentangling the genetic contribution of workers from that of the queen to the colony’s phenotype, as different offspring worker groups from the same queen but a distinct sire cannot be obtained. The mode of reproduction in the honeybee thus condemns a queen to be evaluated with their early mate, be it a PS or SS. This is different from other livestock species, where females can have offspring from several known mates, but even in that case disentangling direct from maternal effects is still difficult [14,15].

Recently, Du et al. [16] explored the effect of data structure on the estimates of genetic parameters in simulated unselected honeybee populations. Among other parameters, they varied the proportion of missing phenotypes, as well as the proportion of controlled and uncontrolled mated queens, with drones always originating from the closed nucleus population. They demonstrated the importance of the proportion of controlled mating for the accuracy of estimates of genetic parameters and obtained unbiased estimates as long as at least 20% of the colonies were recorded. However, they did not explore the impact of SS *vs* PS mating, and the consequences of erroneous assumptions when the true mating strategy is unknown, nor how to model the effect of open mating of DPQ with phenotypic records.

With the increased number of honeybee breeding plans worldwide, records from queens with more diversified mating strategies often need to be considered in genetic analyses. Here we investigate the impact of the mating strategy and the sire modeling on the bias and standard error of genetic parameter estimates and breeding values, using simulation. We focus on datasets with colony phenotypes from both inseminated breeding queens and open-mated DPQs. First, for inseminated breeding queens, we quantify the reduction in the standard error of estimated genetic parameters when the sire was a single DPQ (SS mating) compared to a group of DPQs (PS mating). Second, with the same simulation datasets but assuming that only the dams of DPQ(s) were known in the sire pedigree, we explored how assuming that drones came either from a PS or a SS impacted the genetic evaluation. Lastly, we investigated two alternative strategies to accommodate phenotypic records of open-mated DPQs in the genetic evaluation: either by adding dummy PS in the pedigree, or by adding a non-genetic effect to the evaluation model to account for the origin of the mates of the DPQs.

## Methods

To assess the impact of the mating strategy and the sire modeling on the estimation of genetic parameters and breeding values, we used two simulation sets (Table 1). In both simulation sets, we simulated controlled mating of breeding queens (BQs) (except for initial BQs, from first to third generation, which were open mated) and open mating of DPQs. For all scenarios, 200 replicates were run.

**Table 1:**
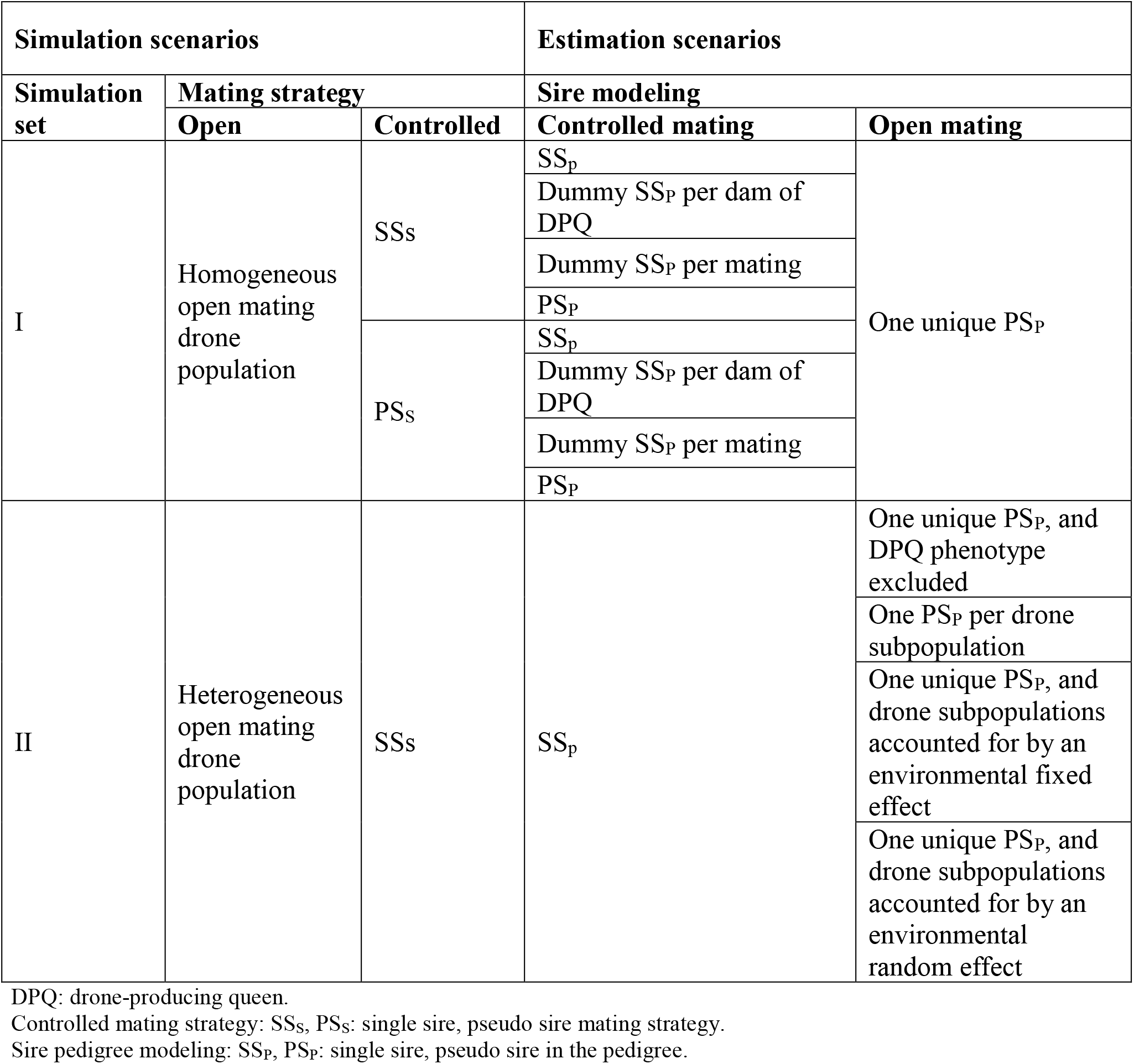
Simulation and estimation scenarios.

| Simulation scenarios |  |  | Estimation scenarios |  |
| --- | --- | --- | --- | --- |
| Simulation set | Mating strategy |  | Sire modeling |  |
|  | Open | Controlled | Controlled mating | Open mating |
| I | Homogeneous open mating drone population | SSs | SS<sub>p</sub> | One unique PS<sub>p</sub> |
|  |  |  | Dummy SS<sub>p</sub> per dam of DPQ |  |
|  |  |  | Dummy SS<sub>p</sub> per mating |  |
|  |  |  | PS<sub>p</sub> |  |
|  |  | PSs | SS<sub>p</sub> |  |
|  |  |  | Dummy SS<sub>p</sub> per dam of DPQ |  |
|  |  |  | Dummy SS<sub>p</sub> per mating |  |
|  |  |  | PS<sub>p</sub> |  |
| II | Heterogeneous open mating drone population | SSs | SS<sub>p</sub> | One unique PS<sub>p</sub>, and DPQ phenotype excluded |
|  |  |  |  | One PS<sub>p</sub> per drone subpopulation |
|  |  |  |  | One unique PS<sub>p</sub>, and drone subpopulations accounted for by an environmental fixed effect |
|  |  |  |  | One unique PS<sub>p</sub>, and drone subpopulations accounted for by an environmental random effect |

DPQ: drone-producing queen.
Controlled mating strategy: SSs, PSs: single sire, pseudo sire mating strategy.
Sire pedigree modeling: SS<sub>p</sub>, PS<sub>p</sub>: single sire, pseudo sire in the pedigree.

### Simulation set I: Sire modeling for controlled mating of breeding queens

In simulation set I, we explored the impact of the sire modeling of controlled mating. In the simulation, we considered two controlled mating strategies for BQs. The first strategy was SS_S_, (the subscript refers to simulation) where a single DPQ was randomly chosen among three sister-DPQs to produce all the drones mating a single BQ. The second strategy was PS_S_, where the three sister-DPQs formed a PS jointly producing the drones mating a single BQ, with random contribution of each sister to the drone pool. In both strategies, the contribution of each group of sister-DPQs to the total drone pool mating all BQs was balanced. All generations of DPQs were open mated to a homogeneous drone population descending from the base population of BQs.

In the estimation, the dam of DPQ(s) that mated BQs was always correctly identified, but four different sire-modeling scenarios were considered to study the consequences of lack of knowledge of whether one or more DPQs were used (Table 1, Fig. 1). First, in case of SS_P_ (where the subscript refers to the way the sire is included in the pedigree), individual DPQs were identified in the pedigree exactly as they were used for mating when SS_S_ was the controlled mating strategy in the simulation; alternatively, one of the three sister-DPQs making up the pseudo sire was randomly identified in the pedigree when pseudo sire mating was used in the simulation (PS_S_). Second, in the ‘dummy SS_P_ per dam of DPQ’ scenario, one single DPQ was identified for all queens mated with drones from the same dam of DPQs. Third, in the ‘dummy SS_P_ per mating’ scenario, a different dummy DPQ was identified in the pedigree for each mating. Lastly, in the PS_P_ scenario, a pseudo sire made up of three sister-DPQs was identified in the pedigree.

**Figure 1:**
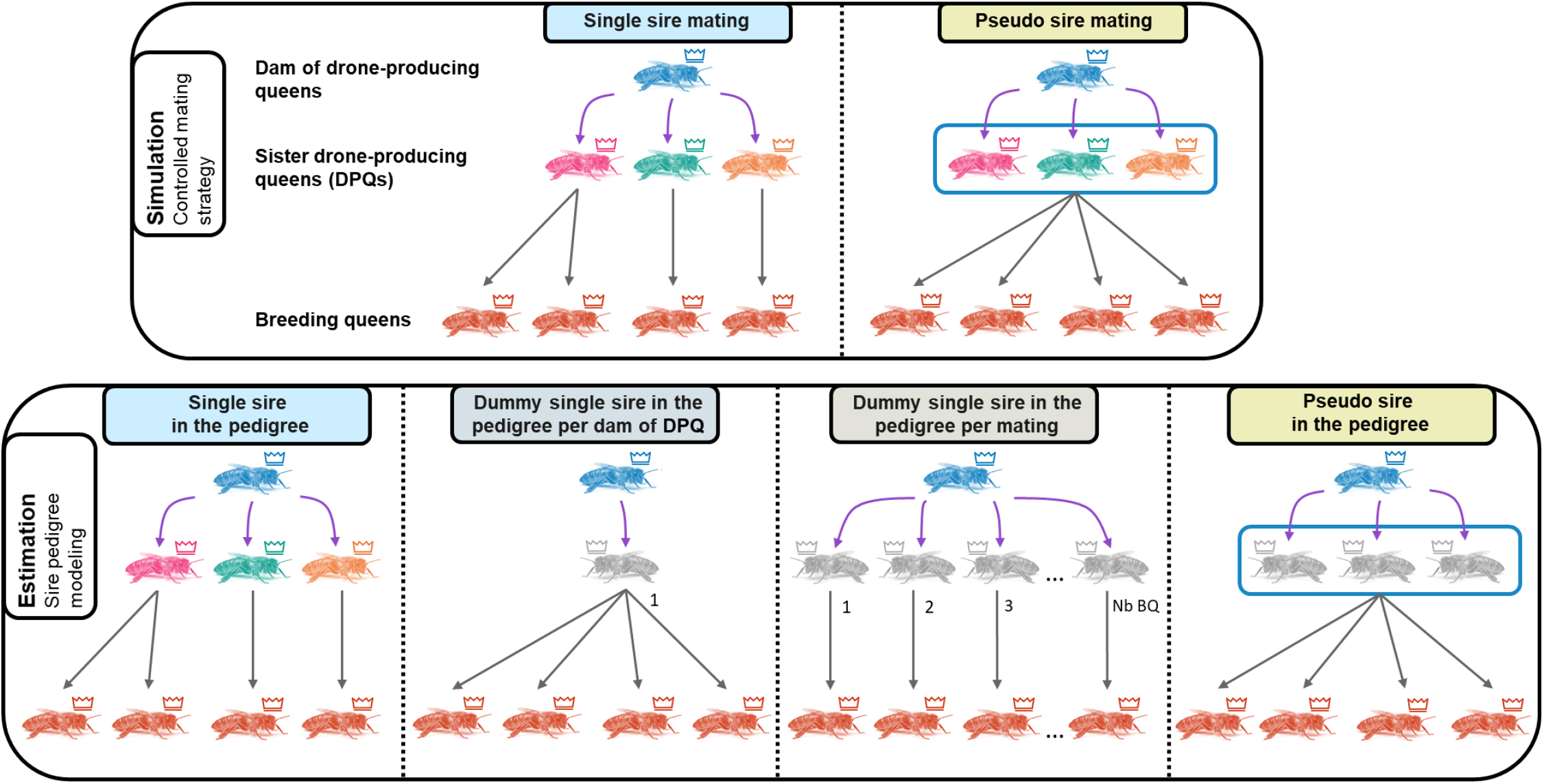
Sire modeling scenarios for controlled mating. In the first set of simulations, either single sire mating (all drones mating a queen are produced by a single drone-producing queen, DPQ) or pseudo sire mating (all drones mating a queen are produced by a group of three sister DPQs) was used. The dam of DPQ(s) was always correctly identified. Irrespective of the true (simulated) mating strategy used, four hypothetical scenarios were used to derive the sire pedigree, to simulate different ways of handling uncertainty in the true mating strategy used. First, in the single sire in the pedigree modeling (SS_P_), individual DPQ were identified exactly as they were used for mating when single sire mating was the controlled mating strategy in the simulation (SS_S_); alternatively, one of the three sister DPQs making up the pseudo sire was randomly identified in the pedigree when pseudo sire mating was used in the simulation (PS_S_). Second, in the ‘dummy SS_P_ per dam of DPQ’ scenario, a unique dummy DPQ was identified for all queens mating with drones from the same dam of DPQs. Third, in the ‘dummy SS_P_ per mating’ scenario, one dummy DPQ was identified in the pedigree for each mating. Lastly, in the pseudo sire in the pedigree modeling (PS_P_), a pseudo sire made of three sister DPQs was identified in the pedigree.

For all four scenarios, the contribution of open mating drones to the colony phenotypes of initial BQs and all DPQs was modeled by genetic effects through the pedigree. One single wild group of a hundred non-inbred and unrelated DPQs (an open mating PS_P_), descending from the base population, was supposed to produce all open mating drones and was uniquely identified in the pedigree files. The open mating PS_P_ was modeled as a group of individuals, and as such had coefficients in the relationship matrix divided by the number of DPQs it was supposed to be composed of (see Additional file 1 for more details).

### Simulation set II: Sire modeling for open mating of drone-producing queens

In simulation set II, we explored the impact of the modeling of the sire of workers of open mated queens. In this simulation set, we only considered SS_S_ mating for BQs, while DPQs were mated to a heterogeneous open mating drone population (in contrast to the homogeneous population in simulation set I). This drone population consisted of two distinct subpopulations with a mean genetic level differing by 3 units, corresponding to approximately one genetic standard deviation of queen effects. Drones from each subpopulation were randomly mated to one half of the DPQ sister-groups.

In the estimation, we appropriately assumed only SS_P_ for BQs, except for initial BQs. For DPQs, we considered four scenarios to model the effect of the subpopulations of drones mating them. First, we eliminated colony phenotypes of DPQs from the performance file altogether, even though selection took place. Second, we accounted for the effect of the drone subpopulations by identifying two dummy open mating PS_P_ in the pedigree as mates of open mated queens: one for each drone subpopulation mating the DPQs. Finally, in the last two scenarios, we accounted for the effect of the drone subpopulations by adding either a fixed or random non-genetic effect in the statistical model describing phenotypes. For the initial BQs in all four scenarios, and also DPQs in the last two scenarios, we identified a unique open mating PS_P_ as mate (as in simulation set I, see Additional file 1).

### Simulation: genetic inheritance, phenotype modeling, and population structure

For both simulation sets I and II, phenotypes of colonies and breeding values of individual queens, drones and worker groups were stochastically simulated [17] based on an infinitesimal model adapted to the honeybee, following Kistler et al. [18].

After the generation of founders, each year, wintering mortality was modeled by randomly eliminating 25% of all BQs and DPQs. Phenotypes were obtained after wintering, within the first year of birth for BQs and within their second year for DPQs. All phenotypes were obtained before selection and reproduction. The resulting generation interval was 1.5 years (one year on the maternal path and two years on the paternal path, Table 2). We simulated ten generations of within-maternal family selection based on phenotypes (one replacement queen per maternal sister-group). On the paternal path, two-thirds of the dam of DPQs were selected each year based on their mean DPQ phenotypes (across-family selection). Three DPQs were then randomly chosen per selected dam family to mate with BQs. Each selected BQ produced 24 BQs and 20 DPQs. After wintering losses, BQ families consisted on average of 18 queens, and DPQ families of 15 queens, which were candidates for selection. Each queen mated with 8 drones, equally contributing to the genetic effect of the worker groups. All colonies performing a same year were affected by a year effect, drawn from a normal distribution centered on zero and with a variance equal to 2/3 of the residual variance.

**Table 2:**
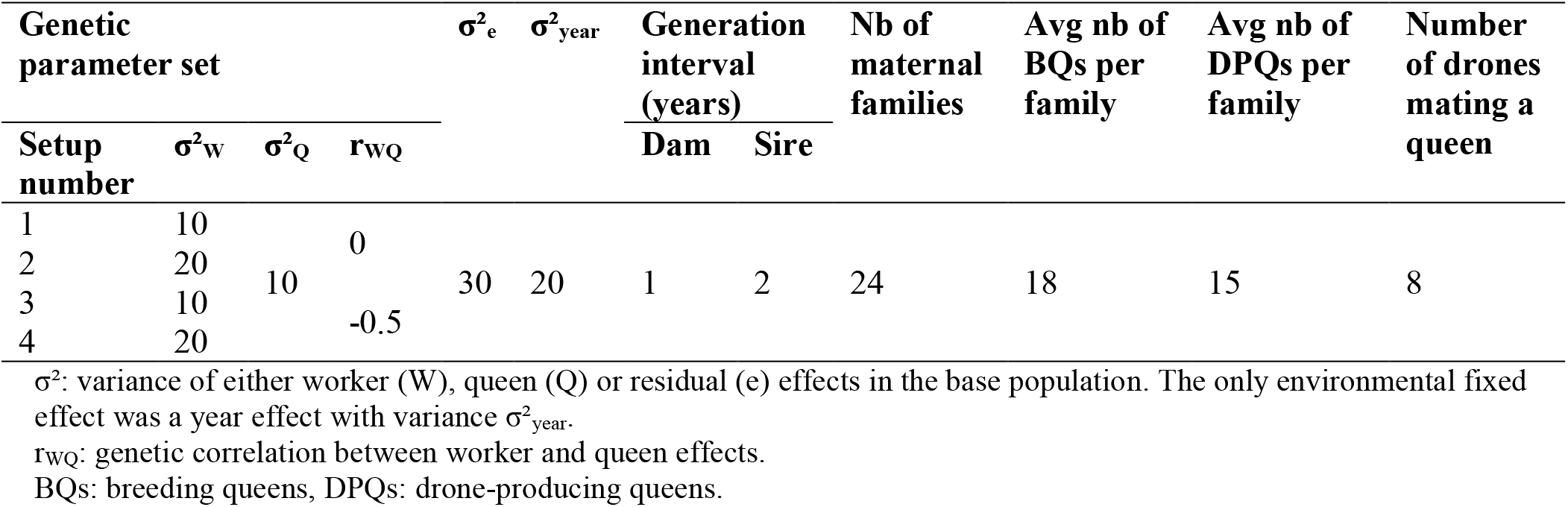
Input parameters for the simulations.

### Simulation: genetic parameter sets

For both simulation sets I and II, four sets of genetic parameters were simulated, following Kistler et al. [18] (Table 2). In scenarios one and three, the variances of worker and queen effects were both equal to a third of the residual variance. In setups two and four, the variance of worker effects was doubled. The genetic correlation between worker and queen effects (r_WQ_) was either null or -0.5. These values represent typical estimates for honeybee production and behavioral traits [19,20,11,21].

### Genetic evaluation: mixed model with queen and worker effects

Pedigree and colony records were used for a single retrospective estimation of genetic parameters and breeding values. The vector of phenotypes **y** was described using a linear mixed model with worker and queen effects [9,10]:

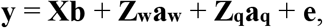

where **b** is the vector of fixed year effects with a corresponding incidence matrix **X, a**_**w**_ the vector of worker effects with incidence matrix **Z**_**w**_, **a**_**q**_ the vector of queen effects with incidence matrix **Z**_**q**_, and **e** the vector of residuals. Breeding values and genetic parameters were estimated using the additive genetic relationships matrix according to Brascamp & Bijma [6,22,23].

For simulation set II, an additional fixed or random effect was added for the subpopulation of the open mating drones for the relevant scenarios.

Genetic parameters were used to obtain BLUP estimated breeding values (EBVs) [24]. The true genetic trend was calculated as the regression coefficient of true breeding values (BVs) of BQs over years, from the fifth generation of the breeding program (when the selection nucleus became closed) to the last tenth generation of selection. The estimated genetic trend was derived similarly based on EBVs instead of BVs.

Starting values used for the AIReML algorithm [25] are shown in Additional file 2: Table S1.

## Results

In all scenarios, at least 99% of the 200 replicates converged, except for simulation set II with equal variance for worker and queen effects, r_WQ_ = -0.5, and a random non-genetic effect for the open mating drone subpopulation, for which 24% of the replicates failed to converge (see Additional file 2: Table S3 and Table S4).

### Results across estimation scenarios

Across all scenarios, no strong biases were observed when the male mate pedigree of BQs was known and properly modeled, and when the effect of drone subpopulations in open mating was accounted for by an environmental effect. In other pedigree modeling scenarios however, the errors on genetic variance components could be large, while errors on 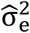 were almost always small. Part of these large errors resulted from under or overestimated variances being compensated by over or underestimated covariances. In addition, there was a tendency to jointly under-(and over-)estimate σ²_W_ and σ²_Q_, particularly for a r_WQ_ of -0.5. Furthermore, within a scenario, the errors on estimated genetic parameters or genetic trends were similar across genetic parameter sets (Except for expected differences, e.g., doubling σ²_W_ reduced the relative errors on 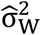). Lastly, errors on 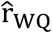 were smaller for r_WQ_ = -0.5 than for r_WQ_ = 0.

### Simulation set I: sire modeling for controlled mating of breeding queens

*Simulate SS or PS controlled mating and estimate genetic parameters accordingly* When the mating of BQs was correctly modeled in the pedigree, no strong biases on (co)variance estimates were observed. However, across scenarios, we still observed a trend to a small underestimation of the variance of queen effects (−2%, Table 3) and a larger one for the genetic trend (Fig. 3) on queen effects (−9% of the true genetic trend, see Additional file 2: Table S2).

**Table 3:** Errors on genetic parameter estimates when male mates for controlled mating are correctly modeled in the pedigree.

| Simulation | Errors on estimated genetic parameters |  |  |  |  |  |  |  |
| --- | --- | --- | --- | --- | --- | --- | --- | --- |
| Controlled mating strategy | $\hat{\sigma}_W^2$ | | | $\hat{\sigma}_Q^2$ | | | $\hat{r}_{WQ}$ | |
|  | Relative bias (%) | Relative SE (%) | Strong deviates % | Relative bias (%) | Relative SE (%) | Strong deviates % | Bias (%) | SE (%) |
| **Genetic parameter set 1 ($\sigma^2_W = 10$, $\sigma^2_Q = 10$, $r_{WQ}=0$)** |  |  |  |  |  |  |  |  |
| SSs | -0.81 | 18.46 | 27 | -3.45 | 17.50 | 27 | 0.011 | 0.152 |
| PSs | 0.20 | 20.79 | 37 | -2.21 | 19.55 | 30 | 0.013 | 0.165 |
| **Genetic parameter set 1 ($\sigma^2_W = 20$, $\sigma^2_Q = 10$, $r_{WQ}=0$)** |  |  |  |  |  |  |  |  |
| SSs | -2.33 | 13.16 | 12 | -3.56 | 17.32 | 30 | 0.017 | 0.130 |
| PSs | -0.31 | 17.42 | 26 | -2.89 | 20.42 | 35 | 0.011 | 0.167 |
| **Genetic parameter set 1 ($\sigma^2_W = 20$, $\sigma^2_Q = 10$, $r_{WQ}=0$)** |  |  |  |  |  |  |  |  |
| SSs | 0.45 | 18.57 | 26 | 0.26 | 15.84 | 18 | -0.017 | 0.099 |
| PSs | -1.01 | 26.13 | 45 | -4.07 | 17.00 | 27 | 0.008 | 0.121 |
| **Genetic parameter set 1 ($\sigma^2_W = 20$, $\sigma^2_Q = 10$, $r_{WQ}=-0.5$)** |  |  |  |  |  |  |  |  |
| SSs | -0.67 | 14.38 | 14 | -0.53 | 18.79 | 30 | -0.002 | 0.092 |
| PSs | -0.38 | 19.75 | 34 | -3.47 | 20.68 | 38 | 0.012 | 0.117 |

$\sigma^2_W$ , $\sigma^2_Q$ , $r_{WQ}$ : genetic variances and correlation for worker and queen effect. Estimates are denoted by $\hat{\cdot}$ ; strong deviates differ by more than 20% from the true values. The relative bias was calculated as $\left( \frac{1}{n_{rep}} \sum_1^{n_{rep}} \left( \frac{\hat{\sigma}_A^2 - \sigma_A^2}{\sigma_A^2} \right) \right)$ . The relative standard error was the standard deviation across replicates of the relative difference between estimates and the true value.
Controlled mating strategy: SSs, PSs: single sire, pseudo sire mating strategy.

**Figure 2:**
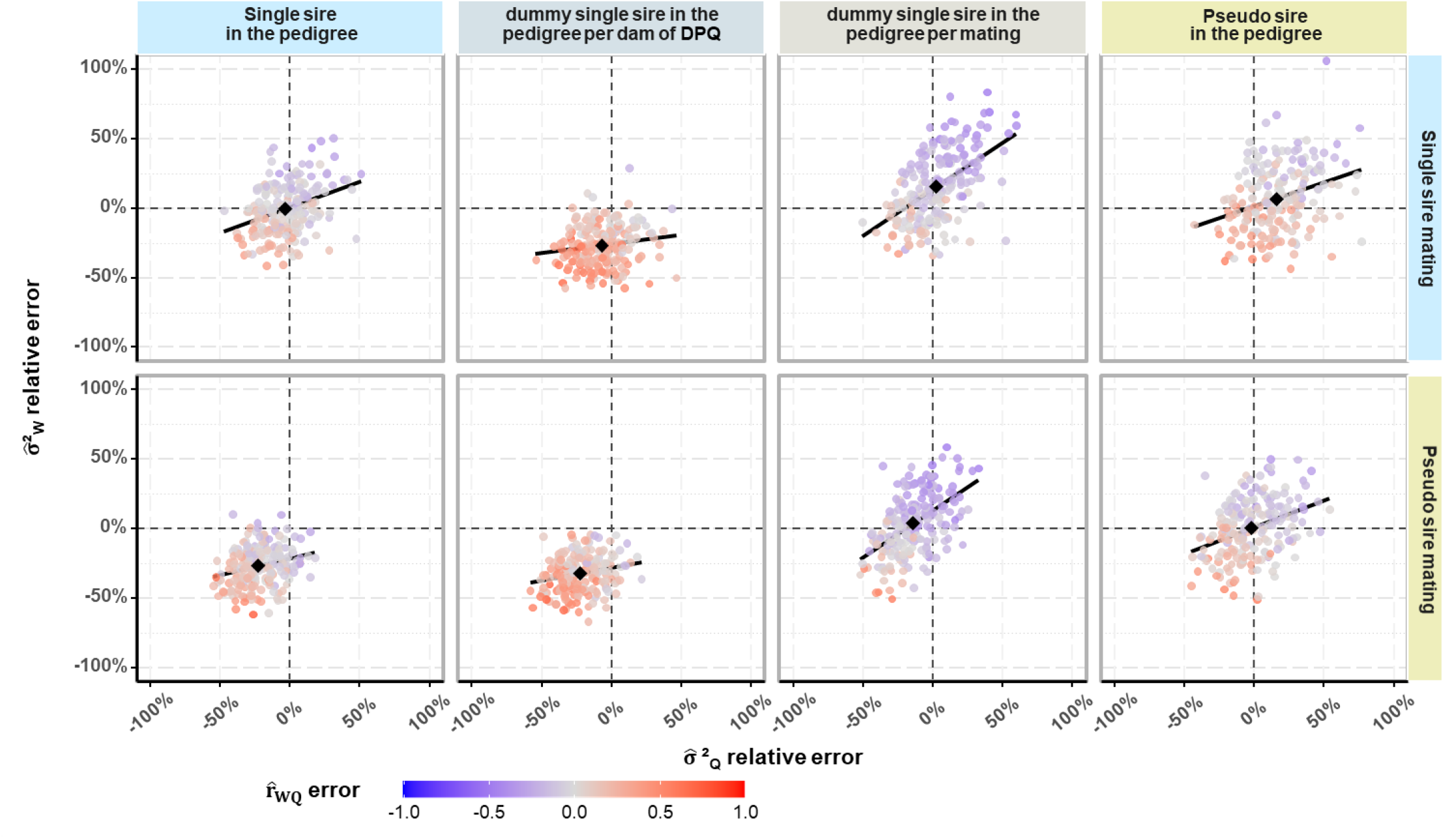
Errors on genetic parameter estimates for sire modeling scenarios for controlled mating and parameter set 1. Black diamond-shaped points indicate the relative bias on genetic variances, black lines the regression lines of relative errors of variance estimates of worker effects on that of queen effects.

**Figure 3:**
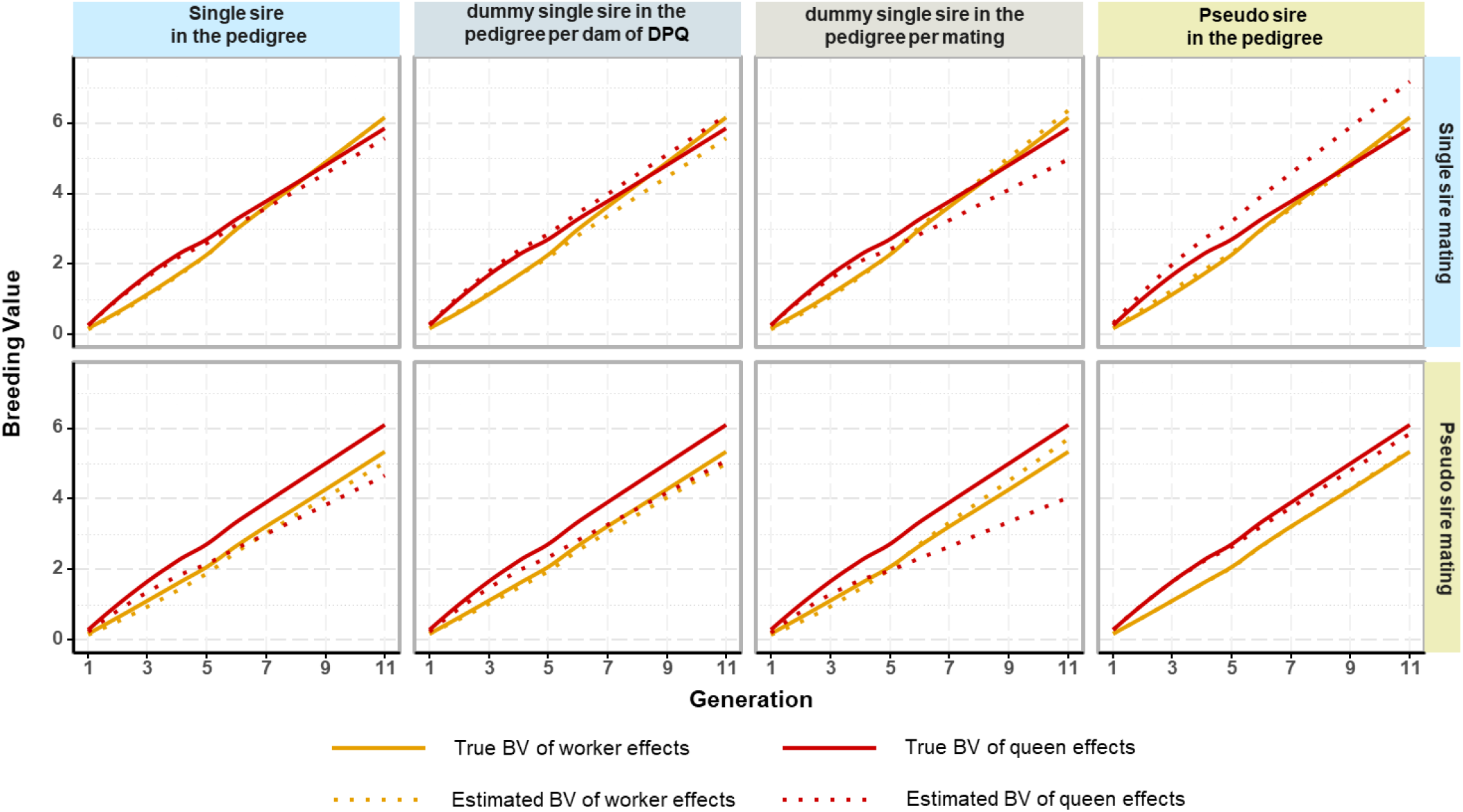
Genetic trends for worker and queen effects for sire modeling scenarios for controlled mating and parameter set 1. From left to right, the results were obtained for i) sire pedigree modeling scenarios with single sire; ii) a dummy single sire per paternal granddam; iii) a dummy single sire for each mating; iv) a pseudo sire.

Across the four genetic parameter sets, about 28% of genetic variance estimates deviated by over 20% from their true values (Table 3). With σ²_W_ = σ²_Q_, in genetic parameter sets 1 and 3, the relative standard error on 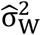 was larger (21%) than on 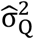 (17%). However, doubling σ²_W_, in genetic parameter sets 2 and 4, decreased the relative standard error of 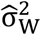 (16%), while that of 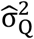 increased slightly (19%).

Compared to PS_S_, for SS_S_ the relative standard error of 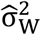 and 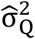 across all genetic parameter sets was reduced in average by 23% and 11%, respectively (Table 3). It also reduced the standard error of the estimated r_WQ_ by 17% on average.

### Simulate SS or PS controlled mating but estimate genetic parameters with incorrect sire modeling alternatives

We report results for parameter set 1 only, as results for other sets are similar. When the dam of DPQs was known, but not if DPQ(s) were used as SS or PS, strong biases could occur in the estimated variance(s), the estimated r_WQ_, or both. Depending on scenarios, the relative bias for genetic variances ranged from -40% to +18%, and the bias for r_WQ_, from -0.19 up to +0.32 (see Table 4 and Additional file 2: Table S3 and Table S4). Likewise for the estimated genetic trend (see Fig. 3 and see Additional file 2: Table S2), wrong mating assumptions led to substantial over or underestimations of the genetic trend on worker and queen effects (from -16% to +33% for BV_W_ and -59% to +44% for BV_Q_).

**Table 4:** Errors on genetic parameter estimates for parameter set 1 when sires for controlled mating are not correctly modeled in the pedigree.

| Sire pedigree modeling<br>for controlled mating | $\hat{\sigma}_W^2$ | | | $\hat{\sigma}_Q^2$ | | | $\hat{r}_{WQ}$ | |
| --- | --- | --- | --- | --- | --- | --- | --- | --- |
|  | Relative<br>bias<br>(%) | Relative<br>SE (%) | Strong<br>deviates<br>% | Relative<br>bias<br>(%) | Relative<br>SE (%) | Strong<br>deviates<br>% | Bias | SE |
| **Controlled mating strategy: SS<sub>s</sub>** |  |  |  |  |  |  |  |  |
| Dummy SS<sub>P</sub> per dam of DPQ | -27.03 | 14.85 | 66 | -6.67 | 17.25 | 28 | 0.248 | 0.174 |
| Dummy SS<sub>P</sub> per mating | 15.08 | 25.38 | 48 | 2.68 | 20.06 | 32 | -0.111 | 0.168 |
| PS<sub>P</sub> | 6.58 | 23.88 | 44 | 16.15 | 21.80 | 42 | 0.069 | 0.169 |
| **Controlled mating strategy: PS<sub>P</sub>** |  |  |  |  |  |  |  |  |
| SS<sub>P</sub> | -26.90 | 14.08 | 68 | -22.71 | 14.93 | 60 | 0.062 | 0.166 |
| Dummy SS<sub>P</sub> per dam of DPQ | -32.27 | 12.82 | 84 | -22.46 | 15.59 | 59 | 0.194 | 0.176 |
| Dummy SS<sub>P</sub> per mating | 3.67 | 22.25 | 38 | -14.02 | 17.32 | 44 | -0.139 | 0.18 |

$\sigma_W^2$ , $\sigma_Q^2$ , $r_{WQ}$ : genetic variances and correlation for worker and queen effect. Estimates are denoted by $\hat{\cdot}$ ; strong deviates differ by more than 20% from the true values. The relative bias was calculated as $\left(\frac{1}{n_{rep}} \sum_1^{n_{rep}} \left(\frac{\hat{\sigma}_A^2 - \sigma_A^2}{\sigma_A^2}\right)\right)$ . The relative standard error was the standard deviation across replicates of the relative difference between estimates and the true value.
SS<sub>s</sub>, PS<sub>s</sub>: single sire, pseudo sire mating strategy; SS<sub>P</sub>, PS<sub>P</sub>: single sire, pseudo sire in the pedigree modeling.

### Simulation set II: sire modeling for open mating of drone-producing queens

In simulation set II, DPQs were mated to a heterogeneous open mating drone population, simulating two distinct subpopulations that differed in their genetic level.

Exclusion of the DPQ records from the genetic evaluation brought no strong biases in the estimated genetic parameters (see Table 5 and Additional file 2: Table S5), even though selection was performed on these colony phenotypes. Nonetheless, across genetic parameter sets, this record’s exclusion increased the standard error of estimated genetic variances (16%), the genetic correlation (30%) and the residual variance (32%), when compared to the scenario including these records and correcting for the drone subpopulations by adding a fixed effect in the evaluation model (see Fig. 4A, Additional file 2: Table S5). Furthermore, exclusion of the DPQ records also led to an underestimation of the genetic trend for queen effects (from -18% to -62%, depending on the genetic parameter set, see Fig. 4B and Additional file 2: Table S6).

**Table 5:** Errors on genetic parameter estimates for parameter set 1 for sire modeling scenarios for open mating.

| $\hat{\sigma}_W^2$ | | | $\hat{\sigma}_Q^2$ | | | $\hat{r}_{WQ}$ | |
| --- | --- | --- | --- | --- | --- | --- | --- |
| Relative bias (%) | Relative SE (%) | Strong deviates % | Relative bias (%) | Relative SE (%) | Strong deviates % | Bias | SE |
| **Modeling one unique open PS<sub>P</sub> and DPQ phenotypes excluded** |  |  |  |  |  |  |  |
| -3.02 | 23.09 | 40 | 1.04 | 21.74 | 34 | -0.021 | 0.195 |
| **Modeling one open PS<sub>P</sub> per drone subpopulation** |  |  |  |  |  |  |  |
| 64.45 | 22.34 | 98 | 21.03 | 20.75 | 54 | -0.245 | 0.110 |
| **Modeling one unique PS<sub>P</sub> and drone subpopulations accounted for by an environmental fixed effect** |  |  |  |  |  |  |  |
| -3.04 | 19.39 | 34 | 0.83 | 18.37 | 24 | 0.003 | 0.146 |
| **Modeling one unique PS<sub>P</sub> and drone subpopulations accounted for by an environmental random effect** |  |  |  |  |  |  |  |
| -2.49 | 18.91 | 33 | 1.24 | 17.84 | 23 | -0.003 | 0.137 |

$\sigma_W^2$ , $\sigma_Q^2$ , $r_{WQ}$ : genetic variances and correlation for worker and queen effect. $\sigma_e^2$ : residual variance. Estimates are denoted by '^'; strong deviates differ by more than 20% from the true values. The relative bias was calculated as $\left( \frac{1}{n_{rep}} \sum_1^{n_{rep}} \left( \frac{\hat{\sigma}_A^2 - \sigma_A^2}{\sigma_A^2} \right) \right)$ . The relative standard error was the standard deviation across replicates of the relative difference between estimates and the true value. The controlled mating strategy was single sire mating (SS<sub>S</sub>).

**Figure 4:**
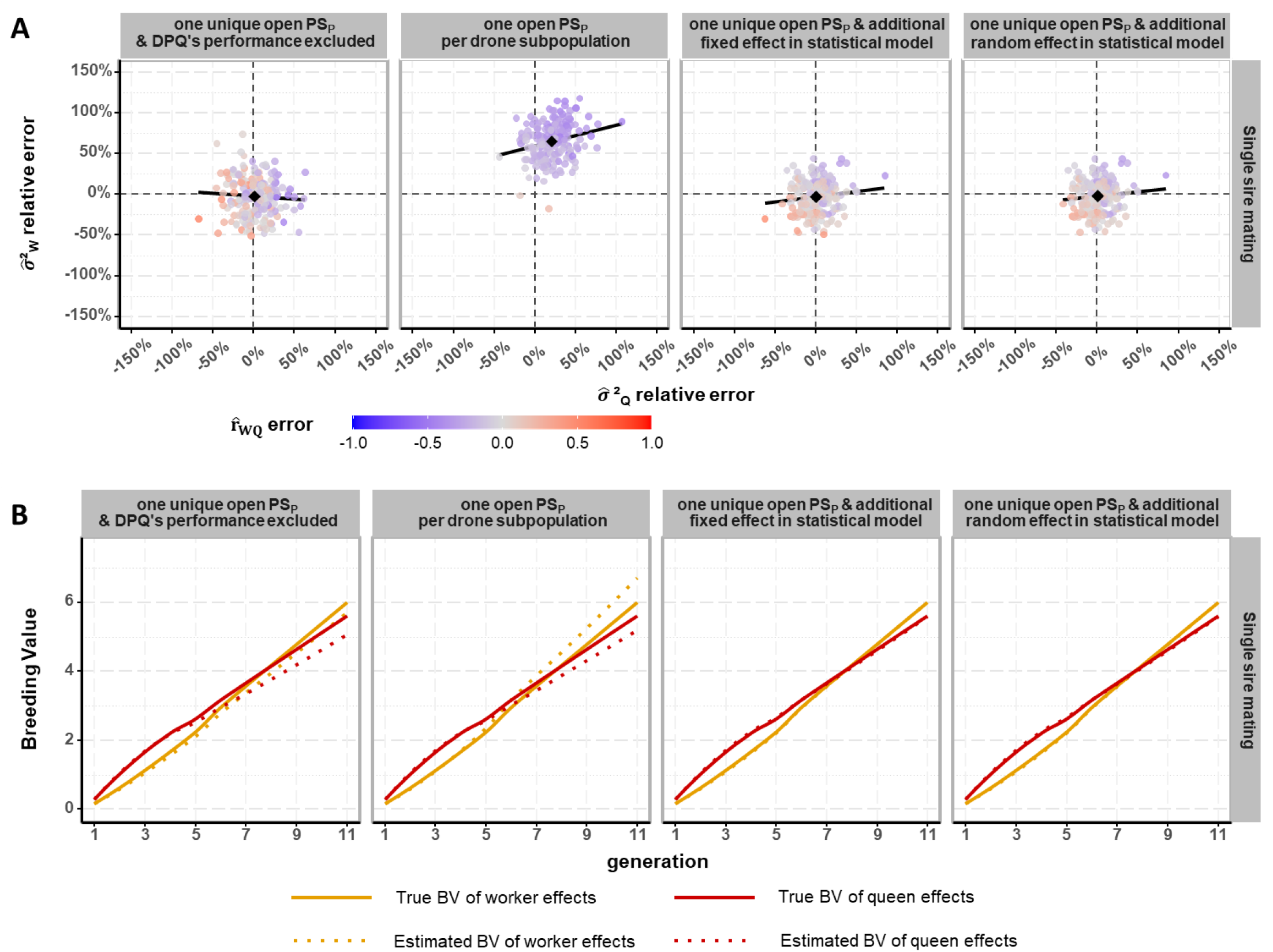
Errors on estimated genetic parameters (A) and breeding values (B), for parameter set 1 and for sire modeling scenarios for open mating. Black diamond-shaped points indicate the relative bias on genetic variances, black lines the regression lines of relative errors of variance estimates of worker effects onto that of queen effects.

Accounting for the open mating of DPQ by modeling a different open mating PS for each drone subpopulation led to a strong overestimation of the variance of worker effects, as well as, to a lesser extent, that of queen effects. With an equal variance for worker and queen effects, bias was around +72% for 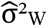 and +26% for 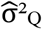, and was halved in scenarios with a doubled variance of worker effects. These overestimated variances were accompanied by an underestimation of the genetic correlation between worker and queen effects of around -0.16 across genetic parameter sets (see Fig. 3A and see Additional file 2: Table S5). In terms of genetic trend, it was overestimated for worker effects (+13% across genetic parameter sets, see Additional file 2: Table S6), while it was underestimated for queen effects by a similar amount as for the scenario with exclusion of the DPQ records from the genetic evaluation.

Finally, the results indicated that the best strategy to account for the open mating drone subpopulations was to add a fixed or random effect in the evaluation model, leading to no or weak biases and the smallest standard errors of genetic parameter and genetic trend estimates. Differences between these two approaches were generally very small, except for the genetic parameter set with equal variance for worker and queen effects and negative r_WQ_. In that case, considering the heterogeneity of the mating of DPQs by a random non-genetic effect impeded convergence in approximately a quarter of the repetitions. When converging, the estimates were very similar to those of the fixed effect model (Fig. 3A).

## Discussion

Here we investigated the impact of the sire modeling on estimates of genetic parameters and genetic trends. First, we studied the controlled mating of BQs, in which we explored the relevance of knowing how the three sister-DPQs of each known dam of DPQs had been used (as SS or PS). Second, we studied the open mating of phenotyped DPQs, accounting for the subpopulations of drones mating these queens in three ways: first, by excluding the DPQ phenotypes; second, through genetic effects by assuming a separate open mating PS for each drone subpopulation; last, by adding an environmental (fixed or random) effect in the evaluation model for each drone subpopulation.

We chose not to vary the number of drones (e.g. from 8 to 16) mating each queen nor the number of selected DPQ per DPQ dam (e.g. from 3 to 8), as preliminary trials showed they impacted only marginally results. Aiming at understanding the impact of erroneous assumptions, we simulated only systematic errors affecting all matings. In real datasets of course, a combination of erroneous and correct assumptions is likely, leading to less extreme results.

### Sire modeling for controlled mating of breeding queen

In breeding plans using isolated mating stations, on each station a dam of DPQs will usually be represented by a group of sister-DPQs. However, in case of instrumental insemination, it is common to use a same dam of DPQs in multiple ways, such as by collecting sperm from a single DPQ or from a few sister-DPQs. Often, however, apart from the dam of DPQs, the usage of DPQs is not recorded. Our results show the importance of knowing as precisely as possible how DPQs of a particular dam were used. If the DPQs were used jointly, as a PS, this should be recorded, as well as the number of sisters composing the group. So far, which DPQs exactly composed the group is not considered in the theory [6], as the sisters making up a PS are supposed unselected, random progeny of their dam. However, knowing this information could be useful when phenotypes of DPQs are used in the breeding value estimation. If DPQs were selected inside a sister-group to form a PS, considering them as random progeny could create biased estimates.

### Most accurate estimates when single sire mating appropriately modeled in the pedigree

Knowing whether SS mating was used is important because the true pedigree relationships between DPQs and descendants are known in that situation, and don’t have to be probabilistically accounted. In addition, the phenotypic information of the single DPQ is better utilized in the variance component and breeding value estimation, further reducing the standard errors of estimates. This gain in precision was notable (Table 3), even though some theoretical flaws existed in the way SS mating was considered in the derivation of the relationship matrix (as noted by Manual Du, personal communication). In fact, for SS mating, we simply used the general formulas of [6], in particular we supposed that the probability of two female offspring to come from a same DPQ (p_2_) followed a Poisson distribution, so that 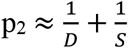, with S the number of sister-DPQs contributing the D drones mating with a queen. When supposing SS mating in the estimation, S equaled 1, leading to p_2_>1.

### Considerable errors when uncertainty in DPQ use in controlled mating

Apart from the loss in precision when SS would erroneously not be assumed, uncertainty in the exact way DPQs were used for mating, and subsequent incorrect assumptions in the genetic evaluation, led to strong biases in both estimated genetic parameters and genetic trends (Table 4).

The observed biases can be synthetized as follows. In one set of scenarios, the genetic relationship between BQs’ offspring and DPQs were overestimated. This occurred when a dummy SS_P_ per dam of DPQs was assumed (when the mating strategy was PS_P_ but also SS_P_). In these scenarios, genetic variances were strongly underestimated, while the genetic correlation was strongly overestimated (Fig. 2). Similarly, when SS_P_ was assumed while the actual mating strategy was PS_S_, also resulting in overestimated genetic relationships between BQs’ offspring and DPQs, led to strongly underestimated variances but affected the genetic correlation much less. In a second set of scenarios, the genetic relationship between BQs’ offspring and DPQs were underestimated. This occurred when a dummy SS_P_ per mating was assumed (whatever the mating strategy). In these scenarios, genetic variance estimates were only moderately biased, but their standard error however became larger, while the genetic correlation was underestimated. Similarly, when PS_P_ was assumed while the mating strategy was SS_S_, also underestimating the genetic relationship between BQs’ offspring and DPQs, led to moderately overestimated variances with larger standard errors, however affecting the genetic correlation only moderately.

### Most accurate estimates for an unknown controlled mating strategy

Not knowing if single sires or pseudo sires were used, modeling a PS_P_ gave the greatest probabilities to get estimates of genetic parameters that do not strongly deviate from their true values (Table 4). Indeed, If the true mating of BQs is unknown, an assumption has to be made on the way DPQs were used in the matings (either as PS_P_ or SS_P_). The most robust approach for estimating genetic parameters was to assume that DPQs were used jointly (PS_P_) to produce the drones that mated a BQ. This gave the most accurate estimates when of course PS_S_ mating was used in the simulation, but also generally when SS_S_ was used, compared to other erroneous assumptions (see Additional file 2: Table S3 and Table S4).

However, assuming PS_P_ when the true mating strategy was unknown was not robust for genetic trend estimation. All assumptions other than SS_P_ for SS_S_ and PS_P_ for PS_S_ led to strong errors in the predicted BVs of either worker or queen effects, or both (Fig. 3 and see Additional file 2: Table S2). Still, when the goal is to minimize the sum of absolute relative errors of the two genetic trends, identifying a dummy SS_P_ per dam of DPQs was the most robust strategy.

### Sire modeling for open mating of drone-producing queens

The best strategy to account for a heterogeneous open mating drone population mating phenotyped DPQs, was to add a non-genetic effect in the evaluation model (Table 5). For open mating, where the drones in each open mating subpopulation have different mean breeding values, excluding the DPQ records from the analysis did not lead to strong biases in the estimated genetic parameters, even though these phenotypes had been used for selection. This held true even if the selection intensity of DPQs was strongly increased (selecting three times less DPQs, data not shown). Adding a dummy open PS_P_ in the pedigree for each of the two drone subpopulations led to overestimated genetic variances. This suggests that the difference of one genetic SD between the two subpopulations ended up in the genetic variance estimate. The difference between the two subpopulations, and consequently between the worker groups they generated, resulted in two sets of DPQ colonies with a systematically different trait value in the data. A difference of one genetic SD between single individuals in the base generation is not unlikely and should not lead to inflated genetic parameter estimates. However, the dummy PS represented a group of 100 individuals, so that the SD of its mean value is much smaller than one genetic SD, as reflected by the small coefficient for the PS on the diagonal of the relationship matrix. Hence, a clear difference between the two PS can only be explained by a large genetic variation in the base generation, resulting in overestimation of the genetic variance.

It appeared that including a non-genetic effect to account for the two subpopulations of drones avoided the biases. Using either a fixed or a random non-genetic effect in the evaluation model brought very similar results. However, we modeled a simplistic situation in which all colonies tested in a year were affected by the same identified environmental effect. Differences between a modeling by fixed or random effects might appear if statistical confounding between the mean genetic effect of open mating drones in an apiary and environmental effects of this apiary exist, in a design where the genetic connectedness between apiaries is limited.

To assess if increasing the genetic variability of open mating drones had an effect on estimated genetic parameters and trends, we increased the (co)variance of worker and queen effects four-fold to generate the BVs of drones, maintaining the difference in the mean genetic value of the two drone subpopulations. However, results (not shown) were very similar to those presented.

### Effect of the breeding nucleus size and genetic correlation between worker and queen effects on estimated genetic parameters and trends

Results (apart from standard errors) of breeding nucleus size on estimated genetic parameters and trends were very similar. To test the need of considering various breeding nucleus sizes, all simulations of the genetic parameter set 1 were repeated using 12 and 36 maternal families. Results were very similar in terms of mean estimated (co)variances, with the expected difference of lower (or higher) standard errors of estimates for bigger (or smaller) nucleus size scenarios. These differences were close to what could be approximated from results obtained with N=24 maternal families by considering the standard errors as proportional to 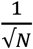.

In addition, these realized standard errors of estimates (calculated from estimated (co)variances over replicates) were in good accordance with mean predicted standard errors by AIReML (see Additional file 2: Table S7).

Last, the standard error of estimated r_WQ_ was lower in genetic parameter sets with a negative r_WQ_ than in those with a null r_WQ_. This was consistent with findings in simulated honeybee datasets [16], and is also in agreement with theoretical predictions [26]. For given genetic variances for worker and queen effects, a negative r_WQ_ also lowers the phenotypic variance, leading to higher heritabilities for the worker and for the queen effect. Another reason for this observation is that we forced the estimated r_WQ_ to be within the parameter space of [-1; 1], resulting in smaller errors for true values nearer to the bounds.

## Conclusion

When breeding queens are mated to drones produced by a single DPQ, as sometimes happens with artificial insemination, and this mating strategy is appropriately modeled, estimates of genetic parameters and genetic trends are more precise as compared to situations where queens are mated to drones produced by a group of sister-DPQs. However, if breeders only record which dam of DPQ(s) is used, but not which particular DPQ or group of sister-DPQs, erroneous assumptions can lead to strong biases in the estimates. When the true mating strategy is unknown, assuming drones come from a group of sister-DPQs leads to the greatest probabilities to get estimates of genetic parameters that do not strongly deviate from their true values. However, strong biases in the estimated genetic trend of queen effects are observed. Moreover, if the DPQs are open mated to a heterogeneous drone population and their phenotypes are used in the genetic analysis, then we recommend adding a non-genetic effect for drone origin in the evaluation model.

## List of abbreviations

BQ: Breeding queen (a selected queen on the dam path)
BLUP: Best linear unbiased predictor
BV: Breeding value
DPQ: Drone-producing queen
EBV: Estimated breeding value
PS: Pseudo sire (a dummy individual representing a group of DPQ used in a mating)
SS: Single sire (a unique DPQ used in a mating)

## Declarations

## Ethics approval and consent to participate

Not applicable

## Consent for publication

Not applicable

## Availability of data and materials

The R simulation code used during the current study is not publicly available while the authors continue to perform active software developments. They may be available from the corresponding author upon reasonable request.

The R code to obtain the inverse of honeybee-specific relationship matrices is available at the AINV-honeybees GitHub repository https://github.com/Tristan-Kistler/AINV-honeybees/. DOI: 10.5281/zenodo.7951334. The version used in this study is v19. Further development will also be available on this repository.

## Authors’ contributions

TK developed the R code for the simulations, performed the simulations and the analysis of the data generated and was involved in the conception of the work, the interpretation of the results and the writing of the original draft. TK modified a FORTRAN program to compute the inverse relationships matrix on the basis of a R code developed and adapted by EWB and a previous FORTRAN program. BB was involved in the conception of the work. EWB, PB and FP supervised the work and were involved in the conception of the work, the methodology choice, the interpretation of the results and revision of the draft. All authors read and approved the final manuscript.

## Acknowledgements

Authors are grateful to Jeremy Vandenplas who wrote a first FORTRAN version of the program to compute the inverse of honeybee relationships matrices.

**Additional files**

**Additional file 1**

Format: Word

Title: Implementation of open mating pseudo sires in the estimation of genetic parameters and breeding values.

**Additional file 2 Table S1**

Format: Word

Title: AIReML starting parameter values and convergence criteria.

The table gives the initial values to estimate (co)variances σ²_W_, σ²_Q_, σ²e and σ_WQ_ respectively for worker, queen and residual effects, and the covariance between worker and queen effects. The true values for genetic variances σ²_W_ and σ²_Q_ were respectively 10 and 20, and for the covariance, respectively 0, -5 or approximately -7, depending on the genetic parameter set. The true σ²_e_ was always equal to 30.

**Additional file 2 Table S2**

Format: Word

Title: True and estimated genetic trends for all genetic parameter sets and sire modeling scenarios for controlled mating.

Description: σ²_W_, r_WQ_: genetic variance of worker effects and genetic correlation between worker and queen effect.

The genetic trends (true and estimate) were calculated as the linear regression coefficients of true breeding values (BV) and estimated breeding values (EBV) for worker (W) and queen (Q) effects over breeding years (after the fifth year of the breeding program, when the nucleus became closed). SS_S,_ PS_S:_ single sire, pseudo sire mating strategy; SS_P,_ PS_P_: single sire, pseudo sire in the pedigree modeling.

**Additional file 2 Table S3**

Format: Word

Title: Errors on estimates for genetic parameter sets with a null r_WQ_ and all sire modeling scenarios for controlled mating.

Description: σ²_W_, σ²_Q_, r_WQ_: genetic variances and correlation for worker and queen effect. σ²_e_: residual variance. Estimates are denoted by ‘^’; strong deviates differ by more than 20% from the true values.

SS_S,_ PS_S:_ single sire, pseudo sire mating strategy; SS_P,_ PS_P_: single sire, pseudo sire in the pedigree modeling.

**Additional file 2 Table S4**

Format: Word

Title: Errors on estimates for genetic parameter sets with a negative r_WQ_ and all sire modeling scenarios for controlled mating.

Description: σ²_W_, σ²_Q_, r_WQ_: genetic variances and correlation for worker and queen effect. σ²_e_: residual variance. Estimates are denoted by ‘^’; strong deviates differ by more than 20% from the true values.

SS_S,_ PS_S:_ single sire, pseudo sire mating strategy; SS_P,_ PS_P_: single sire, pseudo sire in the pedigree modeling.

**Additional file 2 Table S5**

Format: Word

Title: Errors on estimates for all genetic parameter sets and sire modeling scenarios for open mating.

Description: σ²_W_, σ²_Q_, r_WQ_: genetic variances and correlation for worker and queen effect. σ²e: residual variance. Estimates are denoted by ‘^’; strong deviates differ by more than 20% from the true values.

The controlled mating strategy was single sire mating (SS_S_).

**Additional file 2 Table S6**

Format: Word

Title: True and estimated genetic trends for all genetic parameter sets and sire modeling scenarios for open mating.

Description: σ²_W_, r_WQ_: genetic variance of worker effects and genetic correlation between worker and queen effect.

The genetic trends (true and estimate) were calculated as the linear regression coefficients of true breeding values (BV) and estimated breeding values (EBV) for worker (W) and queen (Q) effects over breeding years (from the fifth year of the breeding program, when the nucleus became closed). The controlled mating strategy was single sire mating (SS_S_).

**Additional file 2 Table S7**

Format: Word

Title: AIReML predicted and realized standard errors (SE) of genetic (co)variances.

Description: Predicted SE: mean prediction (by the inverse averaged information matrix) of the SE of genetic (co)variance estimates over repetitions. Realized SE: SD over repetitions of the error on the variance estimates of worker (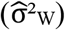) and queen (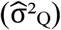) effects, as well as the covariance (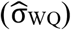). SS_S,_ PS_S:_ single sire, pseudo sire mating strategy; SS_P,_ PS_P_: single sire, pseudo sire in the pedigree modeling.

