## Additional file 1 for "Uncertainty in the mating strategy causes bias and inaccuracy in estimates of genetic parameters in honeybees"

**Implementation of open mating pseudo sires in the estimation of genetic parameters and breeding values**

An open mating pseudo-sire in the pedigree (PS_P_) is a dummy individual considered a founder. It is supposed to be a group of unrelated, non-inbred drone-producing queens (DPQs). It has a coefficient on the diagonal of the (co)variance matrix A that is one over the number (s) of dummy DPQs it is supposed to be made of. Animals (or worker groups) that are paternal offspring of this dummy PS_P_ are only related by 1/(2*s) to this dummy sire.

As an illustration, let’s consider this simplified pedigree:
- BQ1_1* & BQ2_1* are founder breeding queens,
- PS1* & PS2* are founder dummy open mating PS_P_,
- BQ1_2 & BQ2_2 are full-sibs from the mating BQ1_1* × PS1*,
- BQ1_2 & BQ4_2 are maternal half-sibs from the dam BQ2_1*,
- BQ1_2 & BQ3_2 are paternal half-sibs rom the dummy PS1*.

We get the following additive genetic (co)variance matrix:

|  | **BQ1_1*** | **BQ2_1*** | **PS1*** | **PS2*** | **BQ1_2** | **BQ2_2** | **BQ3_2** | **BQ4_2** |
| --- | --- | --- | --- | --- | --- | --- | --- | --- |
| **BQ1_1*** | **1** | 1 | 0 | 0 | 1/2 | 1/2 | 0 | 1/2 |
| **BQ2_1*** | 1 | **1** | 0 | 0 | 0 | 0 | 1/2 | 0 |
| **PS1*** | 0 | 0 | **1/s** | 0 | 1/2s | 1/2s | 1/2s | 0 |
| **PS2*** | 0 | 0 | 0 | **1/s** | 0 | 0 | 0 | 1/2s |
| **BQ1_2** | 1/2 | 0 | 1/2s | 0 | **1** | 1/4 + 1/4s | 1/4s | 1/4 |
| **BQ2_2** | 1/2 | 0 | 1/2s | 0 | 1/4 + 1/4s | **1** | 1/4s | 1/4 |
| **BQ3_2** | 0 | 1/2 | 1/2s | 0 | 1/4s | 1/4s | **1** | 0 |
| **BQ4_2** | 1/2 | 0 | 0 | 1/2s | 1/4 | 1/4 | 0 | **1** |

Founders are distinguished by a ’*’ symbol.
’s’ refers to the number of DPQs making up an open mating PS_P_.

It follows that the additive genetic covariance between:
- full-sibs equals 0.25 + 1/4s,

- maternal half-sibs equals 0.25,

- paternal half-sibs equals 1/4s.

When the number ‘s’ of DPQ constituting the open mating PS_P_ increases, the additive genetic covariance between full-sibs tends towards 0.25, and the one between paternal half-sibs tends towards zero.
