## Additional file 2 for "Uncertainty in the mating strategy causes bias and inaccuracy in estimates of genetic parameters in honeybees"

**Table S1 AIReML starting parameter values and convergence criteria**

| **σ²_W_** | **σ²_Q_** | **σ_WQ_** | **σ²e** | **Converge  criterion** | **Maximum  convergence rounds** | **Variance of the additional  random effect for  open mating drone populations (when used)** | **Total nb of phenotyping records in each genetic analysis** |
| --- | --- | --- | --- | --- | --- | --- | --- |
| 15 | 15 | 0.01 | 20 | 1.00E-12 | 100 | 2 | 8,352 |

The table gives the initial values to estimate (co)variances σ²_W_, σ²_Q_, σ²e and σ_WQ_ respectively for worker, queen and residual effects, and the covariance between worker and queen effects. The true values for genetic variances σ²_W_ and σ²_Q_ were respectively 10 and 20, and for the covariance, respectively 0, -5 or approximately -7, depending on the genetic parameter set. The true σ²_e_ was always equal to 30.

**Table S2** **True and estimated genetic trends for all genetic parameter sets and sire modeling scenarios for controlled mating**

| **Simulation scenario** | | | **Sire pedigree modeling for controlled mating** | **Genetic trend for worker effects** | | **Genetic trend for queen effects** | |
| --- | --- | --- | --- | --- | --- | --- | --- |
| **r_WQ_** | **σ²_W_** | **Controlled mating strategy** |  | **True value** | **Estimate** | **True value** | **Estimate** |
| 0 | 10 | SS_S_ | SS_P_ | 0.62 | 0.62 | 0.50 | 0.47 |
|  |  |  | Dummy SSP per dam of DPQ | 0.62 | 0.54 | 0.50 | 0.53 |
|  |  |  | Dummy SS_P_ per mating | 0.62 | 0.65 | 0.50 | 0.41 |
|  |  |  | PS_P_ | 0.62 | 0.59 | 0.50 | 0.63 |
|  |  | PS_S_ | SS_P_ | 0.52 | 0.50 | 0.54 | 0.40 |
|  |  |  | Dummy SSP per dam of DPQ | 0.52 | 0.49 | 0.54 | 0.43 |
|  |  |  | Dummy SS_P_ per mating | 0.52 | 0.58 | 0.54 | 0.33 |
|  |  |  | PS_P_ | 0.52 | 0.53 | 0.54 | 0.51 |
|  | 20 | SS_S_ | SS_P_ | 1.13 | 1.14 | 0.44 | 0.43 |
|  |  |  | Dummy SSP per dam of DPQ | 1.13 | 1.01 | 0.44 | 0.55 |
|  |  |  | Dummy SS_P_ per mating | 1.13 | 1.18 | 0.44 | 0.34 |
|  |  |  | PS_P_ | 1.13 | 1.09 | 0.44 | 0.61 |
|  |  | PS_S_ | SS_P_ | 0.97 | 0.90 | 0.49 | 0.35 |
|  |  |  | Dummy SSP per dam of DPQ | 0.97 | 0.90 | 0.49 | 0.42 |
|  |  |  | Dummy SS_P_ per mating | 0.97 | 1.04 | 0.49 | 0.25 |
|  |  |  | PS_P_ | 0.97 | 0.97 | 0.49 | 0.46 |
| -1 | 10 | SS_S_ | SS_P_ | 0.38 | 0.40 | 0.24 | 0.21 |
|  |  |  | Dummy SSP per dam of DPQ | 0.38 | 0.32 | 0.24 | 0.27 |
|  |  |  | Dummy SS_P_ per mating | 0.38 | 0.43 | 0.24 | 0.16 |
|  |  |  | PS_P_ | 0.38 | 0.34 | 0.24 | 0.35 |
|  |  | PS_S_ | SS_P_ | 0.27 | 0.28 | 0.33 | 0.21 |
|  |  |  | Dummy SSP per dam of DPQ | 0.27 | 0.26 | 0.33 | 0.22 |
|  |  |  | Dummy SS_P_ per mating | 0.27 | 0.36 | 0.33 | 0.14 |
|  |  |  | PS_P_ | 0.27 | 0.28 | 0.33 | 0.29 |
|  | 20 | SS_S_ | SS_P_ | 0.90 | 0.89 | 0.07 | 0.06 |
|  |  |  | Dummy SSP per dam of DPQ | 0.90 | 0.76 | 0.07 | 0.17 |
|  |  |  | Dummy SS_P_ per mating | 0.90 | 0.94 | 0.07 | -0.01 |
|  |  |  | PS_P_ | 0.90 | 0.82 | 0.07 | 0.21 |
|  |  | PS_S_ | SS_P_ | 0.69 | 0.67 | 0.18 | 0.10 |
|  |  |  | Dummy SSP per dam of DPQ | 0.69 | 0.66 | 0.18 | 0.12 |
|  |  |  | Dummy SS_P_ per mating | 0.69 | 0.80 | 0.18 | -0.01 |
|  |  |  | PS_P_ | 0.69 | 0.70 | 0.18 | 0.16 |

σ²_W_, r_WQ_: genetic variance of worker effects and genetic correlation between worker and queen effect.
The annual genetic trends (true and estimate) were calculated as the linear regression coefficients of true breeding values (BV) and estimated breeding values (EBV) for worker (W) and queen (Q) effects over breeding years (after the fifth yearof the breeding program, when the nucleus became closed).
SS_S,_ PS_S:_ single sire, pseudo sire mating strategy; SS_P,_ PS_P_: single sire, pseudo sire in the pedigree modeling.

**Table S3 Errors on estimates for genetic parameter sets with a null r_WQ_ and all sire modeling scenarios for controlled mating**

| **Simulation scenario** | **Estimation scenario** | **Estimates** | | | | | | | | | |
| --- | --- | --- | --- | --- | --- | --- | --- | --- | --- | --- | --- |
|  |  | ${\hat{\boldsymbol{\sigma}}}_{\mathbf{e}}^{\mathbf{2}}$ | | ${\hat{\boldsymbol{\sigma}}}_{\mathbf{W}}^{\mathbf{2}}$ | | | ${\hat{\boldsymbol{\sigma}}}_{\mathbf{WQ}}^{\mathbf{2}}$ | | | ${\hat{\mathbf{r}}}_{\mathbf{WQ}}$ | |
|  |  | **Relative bias (%)** | **Relative SE (%)** | **Relative bias (%)** | **Relative SE (%)** | **strong deviates (%)** | **Relative bias (%)** | **Relative SE (%)** | **strong deviates (%)** | **Bias** | **SE** |
| **Genetic parameter set 1 (σ²_W_ = 10, σ²_Q_ = 10, r_WQ_=0)** | | | | | | | | | | | |
| SS_S_ | SS_P_ | 1.08 | 3.47 | -0.81 | 18.46 | 27 | -3.45 | 17.50 | 27 | 0.011 | 0.152 |
|  | Dummy SS_P_ per dam of DPQ | 1.18 | 3.71 | -27.03 | 14.85 | 66 | -6.67 | 17.25 | 28 | 0.248 | 0.174 |
|  | Dummy SS_P_ per mating | 0.88 | 3.60 | 15.08 | 25.38 | 48 | 2.68 | 20.06 | 32 | -0.111 | 0.168 |
|  | PS_P_ | -5.97 | 4.52 | 6.58 | 23.88 | 44 | 16.15 | 21.80 | 42 | 0.069 | 0.169 |
| PS_S_ | SS_P_ | 7.86 | 3.12 | -26.90 | 14.08 | 68 | -22.71 | 14.93 | 60 | 0.062 | 0.166 |
|  | Dummy SS_P_ per dam of DPQ | 6.65 | 3.34 | -32.27 | 12.82 | 84 | -22.46 | 15.59 | 59 | 0.194 | 0.176 |
|  | Dummy SS_P_ per mating | 6.31 | 3.23 | 3.67 | 22.25 | 38 | -14.02 | 17.32 | 44 | -0.139 | 0.180 |
|  | PS_P_ | 0.58 | 4.11 | 0.20 | 20.79 | 37 | -2.21 | 19.55 | 30 | 0.013 | 0.165 |
| **Genetic parameter set 1 (σ²_W_ = 20, σ²_Q_ = 10, r_WQ_=0)** | | | | | | | | | | | |
| SS_S_ | SS_P_ | 1.10 | 4.09 | -2.33 | 13.16 | 12 | -3.56 | 17.32 | 30 | 0.017 | 0.130 |
|  | Dummy SS_P_ per dam of DPQ | 1.75 | 4.40 | -24.10 | 12.19 | 64 | -12.49 | 17.01 | 34 | 0.324 | 0.147 |
|  | Dummy SS_P_ per mating | 0.68 | 4.22 | 10.20 | 17.85 | 30 | 7.66 | 19.36 | 28 | -0.135 | 0.150 |
|  | PS_P_ | -6.70 | 5.39 | 4.78 | 17.93 | 28 | 13.48 | 21.36 | 45 | 0.105 | 0.158 |
| PS_S_ | SS_P_ | 9.90 | 3.92 | -29.06 | 11.07 | 80 | -25.03 | 15.40 | 62 | 0.084 | 0.162 |
|  | Dummy SS_P_ per dam of DPQ | 7.73 | 4.42 | -29.95 | 11.49 | 80 | -27.50 | 15.64 | 68 | 0.255 | 0.179 |
|  | Dummy SS_P_ per mating | 6.69 | 4.04 | -1.32 | 15.17 | 19 | -7.51 | 18.47 | 36 | -0.185 | 0.153 |
|  | PS_P_ | 0.70 | 5.28 | -0.31 | 17.42 | 26 | -2.89 | 20.42 | 35 | 0.011 | 0.167 |

σ²_W_, σ²_Q_, r_WQ_: genetic variances and correlation for worker and queen effect. σ²_e_: residual variance. Estimates are denoted by ’^’; strong deviates differ by more than 20% from the true values.
SS_S,_ PS_S:_ single sire, pseudo sire mating strategy; SS_P,_ PS_P_: single sire, pseudo sire in the pedigree modeling.

**Table S4 Errors on estimates for genetic parameter sets with a negative r_WQ_ and all sire modeling scenarios for controlled mating**

| **Simulation scenario** | **Estimation scenario** | **Estimates** | | | | | | | | | |
| --- | --- | --- | --- | --- | --- | --- | --- | --- | --- | --- | --- |
|  |  | ${\hat{\boldsymbol{\sigma}}}_{\mathbf{e}}^{\mathbf{2}}$ | | ${\hat{\boldsymbol{\sigma}}}_{\mathbf{W}}^{\mathbf{2}}$ | | | ${\hat{\boldsymbol{\sigma}}}_{\mathbf{WQ}}^{\mathbf{2}}$ | | | ${\hat{\mathbf{r}}}_{\mathbf{WQ}}$ | |
|  |  | **Relative bias (%)** | **Relative SE (%)** | **Relative bias (%)** | **Relative SE (%)** | **strong deviates (%)** | **Relative bias (%)** | **Relative SE (%)** | **strong deviates (%)** | **Bias** | **SE** |
| **Genetic parameter set 1 (σ²_W_ = 10, σ²_Q_ = 10, r_WQ_=-0.5)** | | | | | | | | | | | |
| SS_S_ | SS_S_ | 0.28 | 2.81 | 0.45 | 18.57 | 26 | 0.26 | 15.84 | 18 | -0.017 | 0.099 |
|  | Dummy SS_P_ per dam of DPQ | 1.65 | 2.88 | -33.35 | 16.55 | 81 | -12.98 | 15.58 | 34 | 0.156 | 0.142 |
|  | Dummy SS_P_ per mating | -0.32 | 2.94 | 17.89 | 28.17 | 55 | 7.09 | 18.70 | 28 | -0.074 | 0.107 |
|  | PS_P_ | -3.16 | 3.42 | 9.32 | 27.67 | 44 | 10.31 | 19.34 | 33 | 0.054 | 0.118 |
| PS_S_ | PS_S_ | 5.25 | 2.34 | -36.10 | 16.38 | 82 | -25.89 | 13.01 | 68 | 0.063 | 0.136 |
|  | Dummy SS_P_ per dam of DPQ | 4.89 | 2.37 | -40.04 | 15.47 | 90 | -24.79 | 14.15 | 62 | 0.105 | 0.146 |
|  | Dummy SS_P_ per mating | 3.40 | 2.39 | -1.78 | 26.23 | 49 | -10.57 | 16.33 | 34 | -0.078 | 0.116 |
|  | PS_P_ | 0.90 | 2.77 | -1.01 | 26.13 | 45 | -4.07 | 17.00 | 27 | 0.008 | 0.121 |
| **Genetic parameter set 1 (σ²_W_ = 20, σ²_Q_ = 10, r_WQ_=-0.5)** | | | | | | | | | | | |
| SS_S_ | SS_S_ | 0.03 | 3.07 | -0.67 | 14.38 | 14 | -0.53 | 18.79 | 30 | -0.002 | 0.092 |
|  | Dummy SS_P_ per dam of DPQ | 2.62 | 3.21 | -29.96 | 12.27 | 76 | -24.95 | 17.14 | 64 | 0.259 | 0.157 |
|  | Dummy SS_P_ per mating | -1.04 | 3.28 | 10.88 | 19 | 34 | 11.38 | 22.72 | 44 | -0.074 | 0.096 |
|  | PS_P_ | -2.77 | 3.84 | 8.41 | 19.69 | 34 | 4.40 | 22.82 | 39 | 0.082 | 0.127 |
| PS_S_ | PS_S_ | 5.85 | 2.72 | -37.40 | 11.58 | 94 | -29.17 | 14.24 | 74 | 0.103 | 0.122 |
|  | Dummy SS_P_ per dam of DPQ | 5.36 | 2.83 | -37.16 | 12.04 | 92 | -31.37 | 15.81 | 78 | 0.176 | 0.147 |
|  | Dummy SS_P_ per mating | 2.72 | 2.78 | -8.54 | 16.23 | 29 | -6.96 | 18.43 | 34 | -0.076 | 0.099 |
|  | PS_P_ | 0.70 | 3.30 | -0.38 | 19.75 | 34 | -3.47 | 20.68 | 38 | 0.012 | 0.117 |

σ²_W_, σ²_Q_, r_WQ_: genetic variances and correlation for worker and queen effect. σ²_e_: residual variance. Estimates are denoted by ’^’; strong deviates differ by more than 20% from the true values.
SS_S,_ PS_S:_ single sire, pseudo sire mating strategy; SS_P,_ PS_P_: single sire, pseudo sire in the pedigree modeling.

**Table S5** **Errors on estimates for all genetic parameter sets and sire modeling scenarios for open mating**

| **Simulation scenario** | | **Estimation scenario** | **Estimates** | | | | | | | | | | |
| --- | --- | --- | --- | --- | --- | --- | --- | --- | --- | --- | --- | --- | --- |
| **r_dm_** | **σ²_W_** | **Sire pedigree modeling for open mating** | **convergence (%)** | ${\hat{\boldsymbol{\sigma}}}_{\mathbf{e}}^{\mathbf{2}}$ | | ${\hat{\boldsymbol{\sigma}}}_{\mathbf{W}}^{\mathbf{2}}$ | | | ${\hat{\boldsymbol{\sigma}}}_{\mathbf{WQ}}^{\mathbf{2}}$ | | | ${\hat{\mathbf{r}}}_{\mathbf{WQ}}$ | |
|  |  |  |  | **Relative bias (%)** | **Relative SE (%)** | **Relative bias (%)** | **Relative SE (%)** | **% strong deviates** | **Relative bias (%)** | **Relative SE (%)** | **% strong deviates** | **Bias** | **SE** |
| 0 | 10 | One unique open PS_P_,  and DPQ performance excluded | 100 | 0.80 | 5.09 | -3.02 | 23.09 | 40 | 1.04 | 21.74 | 34 | -0.021 | 0.195 |
|  |  | One open PS_P_ per drone subpopulation | 100 | -1.31 | 3.87 | 64.45 | 22.34 | 98 | 21.03 | 20.75 | 54 | -0.245 | 0.110 |
|  |  | One unique PSP, and drone subpopulations accounted for by an environmental fixed effect | 100 | 0.51 | 3.82 | -3.04 | 19.39 | 34 | 0.83 | 18.37 | 24 | 0.003 | 0.146 |
|  |  | One unique PSP, and drone subpopulations accounted for by an environmental random effect | 98 | 0.44 | 3.75 | -2.49 | 18.91 | 33 | 1.24 | 17.84 | 23 | -0.003 | 0.137 |
|  | 20 | One unique open PS_P_,  and DPQ performance excluded | 100 | 0.80 | 5.52 | 0.48 | 16.03 | 21 | -2.20 | 25.70 | 47 | -0.013 | 0.189 |
|  |  | One open PS_P_ per drone subpopulation | 100 | -0.12 | 4.15 | 28.86 | 14.14 | 68 | 6.99 | 21.82 | 38 | -0.149 | 0.130 |
|  |  | One unique PSP, and drone subpopulations accounted for by an environmental fixed effect | 100 | 1.02 | 4.24 | -0.30 | 13.77 | 14 | -3.99 | 20.11 | 32 | 0.004 | 0.148 |
|  |  | One unique PSP, and drone subpopulations accounted for by an environmental random effect | 100 | 1.03 | 4.23 | -0.33 | 13.75 | 14 | -4.01 | 20.10 | 32 | 0.004 | 0.148 |
| -0.5 | 10 | One unique open PS_P_,  and DPQ performance excluded | 100 | 0.26 | 3.55 | 0.75 | 22.54 | 36 | -2.00 | 20.58 | 36 | 0.001 | 0.147 |
|  |  | One open PS_P_ per drone subpopulation | 100 | -1.81 | 2.94 | 78.66 | 24.48 | 100 | 31.00 | 20.63 | 74 | -0.145 | 0.073 |
|  |  | One unique PSP, and drone subpopulations accounted for by an environmental fixed effect | 100 | 0.38 | 2.86 | 0.03 | 21.15 | 32 | -1.14 | 17.67 | 27 | 0.005 | 0.115 |
|  |  | One unique PSP, and drone subpopulations accounted for by an environmental random effect | 76 | 0.10 | 2.76 | 5.69 | 19.57 | 29 | -0.83 | 18.17 | 28 | 0.016 | 0.116 |
|  | 20 | One unique open PS_P_,  and DPQ performance excluded | 100 | 0.54 | 4.20 | -1.52 | 16.47 | 20 | -2.31 | 22.09 | 36 | -0.014 | 0.132 |
|  |  | One open PS_P_ per drone subpopulation | 100 | -0.96 | 2.98 | 32.36 | 15.08 | 78 | 17.77 | 21.08 | 46 | -0.099 | 0.078 |
|  |  | One unique PSP, and drone subpopulations accounted for by an environmental fixed effect | 100 | 0.65 | 3.04 | -2.13 | 14.68 | 15 | -3.21 | 19.13 | 34 | 0.003 | 0.102 |
|  |  | One unique PSP, and drone subpopulations accounted for by an environmental random effect | 100 | 0.65 | 3.04 | -2.14 | 14.67 | 15 | -3.21 | 19.11 | 34 | 0.003 | 0.102 |

σ²_W_, σ²_Q_, r_WQ_: genetic variances and correlation for worker and queen effect. σ²_e_: residual variance. Estimates are denoted by ’^’; strong deviates differ by more than 20% from the true values.
The controlled mating strategy was single sire mating (SS_S_).

**Table S6** **True and estimated genetic trends for all genetic parameter sets and sire modeling scenarios for open mating**

| **r_WQ_** | **σ²_W_** | **Sire pedigree modeling for open mating** | **Genetic trend for worker effects** | | **Genetic trend for queen effects** | |
| --- | --- | --- | --- | --- | --- | --- |
|  |  |  | **True value** | **Estimate** | **True value** | **Estimate** |
| 0 | 10 | one unique open PS_P_,  and DPQ performance excluded | 0.59 | 0.57 | 0.47 | 0.40 |
|  |  | one open PS_P_ per drone subpopulation | 0.59 | 0.69 | 0.47 | 0.41 |
|  |  | one unique PSP, and drone subpopulations accounted for by an environmental fixed effect | 0.59 | 0.60 | 0.47 | 0.46 |
|  |  | one unique PSP, and drone subpopulations accounted for by an environmental random effect | 0.59 | 0.60 | 0.48 | 0.46 |
|  | 20 | one unique open PS_P_,  and DPQ performance excluded | 1.14 | 1.10 | 0.42 | 0.35 |
|  |  | one open PS_P_ per drone subpopulation | 1.14 | 1.21 | 0.42 | 0.36 |
|  |  | one unique PSP, and drone subpopulations accounted for by an environmental fixed effect | 1.14 | 1.13 | 0.42 | 0.41 |
|  |  | one unique PSP, and drone subpopulations accounted for by an environmental random effect | 1.14 | 1.13 | 0.42 | 0.41 |
| -0.5 | 10 | one unique open PS_P_,  and DPQ performance excluded | 0.40 | 0.41 | 0.21 | 0.13 |
|  |  | one open PS_P_ per drone subpopulation | 0.40 | 0.49 | 0.21 | 0.15 |
|  |  | one unique PSP, and drone subpopulations accounted for by an environmental fixed effect | 0.40 | 0.41 | 0.21 | 0.18 |
|  |  | one unique PSP, and drone subpopulations accounted for by an environmental random effect | 0.42 | 0.47 | 0.20 | 0.18 |
|  | 20 | one unique open PS_P_,  and DPQ performance excluded | 0.88 | 0.87 | 0.07 | -0.01 |
|  |  | one open PS_P_ per drone subpopulation | 0.88 | 0.95 | 0.07 | 0.01 |
|  |  | one unique PSP, and drone subpopulations accounted for by an environmental fixed effect | 0.88 | 0.88 | 0.07 | 0.04 |
|  |  | one unique PSP, and drone subpopulations accounted for by an environmental random effect | 0.88 | 0.88 | 0.07 | 0.05 |

σ²_W_, r_WQ_: genetic variance of worker effects and genetic correlation between worker and queen effect.
The genetic trends (true and estimate) were calculated as the linear regression coefficients of true breeding values (BV) and estimated breeding values (EBV) for worker (W) and queen (Q) effects over breeding years (from the fifth year of the breeding program, when the nucleus became closed).
The controlled mating strategy was single sire mating (SS_S_).

**Table S7** **AIReML predicted and realized standard errors (SE) of genetic (co)variances**

| **Controlled mating  strategy** | **Sire pedigree modeling for controlled mating** | **SE(σ²_W_)** | | **SE(σ²_Q_)** | | **SE(σ_WQ_)** | |
| --- | --- | --- | --- | --- | --- | --- | --- |
|  |  | **Predicted** | **Realized** | **Predicted** | **Realized** | **Predicted** | **Realized** |
| **Nucleus size: 12 Breeding Queens** | | | | | | | |
| SS_S_ | SS_P_ | 2.75 | 2.65 | 2.44 | 2.56 | 2.07 | 1.93 |
|  | dummy SS_P_ per dam of DPQ | 2.25 | 2.18 | 2.43 | 2.57 | 1.89 | 1.75 |
|  | dummy SS_P_ per mating | 3.72 | 3.71 | 2.75 | 2.79 | 2.76 | 2.78 |
|  | PS_P_ | 3.49 | 3.33 | 3.04 | 3.23 | 2.56 | 2.35 |
| PS_S_ | SS_P_ | 2.28 | 2.28 | 2.07 | 1.99 | 1.75 | 1.71 |
|  | dummy SSP per dam of DPQ | 2.20 | 2.13 | 2.21 | 2.14 | 1.80 | 1.84 |
|  | dummy SS_P_ per mating | 3.45 | 3.41 | 2.44 | 2.50 | 2.51 | 2.71 |
|  | PS_P_ | 3.43 | 3.44 | 2.75 | 2.79 | 2.47 | 2.53 |
| **Nucleus size: 24 Breeding Queens** | | | | | | | |
| SS_S_ | SS_P_ | 1.91 | 1.85 | 1.68 | 1.75 | 1.42 | 1.47 |
|  | dummy SSP per dam of DPQ | 1.58 | 1.49 | 1.67 | 1.73 | 1.31 | 1.27 |
|  | dummy SS_P_ per mating | 2.58 | 2.54 | 1.91 | 2.01 | 1.91 | 2.05 |
|  | PS_P_ | 2.46 | 2.39 | 2.11 | 2.18 | 1.80 | 1.79 |
| PS_S_ | SS_P_ | 1.61 | 1.41 | 1.46 | 1.49 | 1.24 | 1.16 |
|  | dummy SSP per dam of DPQ | 1.54 | 1.28 | 1.51 | 1.56 | 1.24 | 1.13 |
|  | dummy SS_P_ per mating | 2.46 | 2.23 | 1.72 | 1.73 | 1.78 | 1.77 |
|  | PS_P_ | 2.41 | 2.08 | 1.92 | 1.96 | 1.73 | 1.54 |
| **Nucleus size: 36 Breeding Queens** | | | | | | | |
| SS_S_ | SS_P_ | 1.55 | 1.67 | 1.36 | 1.34 | 1.16 | 1.27 |
|  | dummy SSP per dam of DPQ | 1.27 | 1.33 | 1.34 | 1.34 | 1.05 | 1.09 |
|  | dummy SS_P_ per mating | 2.09 | 2.21 | 1.54 | 1.54 | 1.55 | 1.68 |
|  | PS_P_ | 1.97 | 2.12 | 1.70 | 1.72 | 1.44 | 1.52 |
| PS_S_ | SS_P_ | 1.30 | 1.21 | 1.17 | 1.22 | 0.99 | 0.93 |
|  | dummy SSP per dam of DPQ | 1.23 | 1.13 | 1.20 | 1.25 | 0.98 | 0.93 |
|  | dummy SS_P_ per mating | 1.97 | 1.82 | 1.38 | 1.40 | 1.42 | 1.38 |
|  | PS_P_ | 1.93 | 1.80 | 1.53 | 1.59 | 1.37 | 1.30 |

Predicted SE: mean prediction (by the inverse averaged information matrix) of the SE of genetic (co)variance estimates over repetitions. Realized SE: SD over repetitions of the error on the variance estimates of worker ($\hat{\sigma}$²_W_) and queen ($\hat{\sigma}$²_Q_) effects, as well as the covariance ($\hat{\sigma}$_WQ_).
SS_S,_ PS_S:_ single sire, pseudo sire mating strategy; SS_P,_ PS_P_: single sire, pseudo sire in the pedigree modeling.
